# How Does White Matter Registration Affect Tractography Alignment?

**DOI:** 10.1101/2022.10.27.513915

**Authors:** Gabriele Amorosino, Emanuele Olivetti, Jorge Jovicich, Paolo Avesani

## Abstract

Tractography is a powerful method to represent the structural connectivity of the brain white matter. Nevertheless, the comparison of these data structures between two individuals is still an open challenge because of their complexity, e.g. digital representation of millions of fibers as polylines. The scientific community spent a meaningful effort to develop new methods of white matter registration aiming to take advantage of diffusion MRI models. Despite the effort to improve the registration of the white matter, little is known about the effect of the registration on tractogram alignment. The main issue for an empirical evaluation is the lack of ground truth, e.g. a sample of data where the correct alignment is validated by experts. This work aims to overcome this drawback by proposing an evaluation framework based on the matching of homologous fiber structures, e.g. known neuroanatomical bundles. The contribution is a quantitative comparison of how the most representative methods of white matter registration affect tractogram alignment.

## 1. INTRODUCTION

Tractography is a data derivative that encodes the fiber pathways of brain white matter. This type of data results from a composite processing pipeline: (i) Diffusion-weighted magnetic resonance imaging (DWI) acquisition, (ii) DWI preprocessing, (iii) diffusion model reconstruction, and (iv) fiber tracking [1]. The output is a tractogram, a collection of 10^5*−*6^ streamlines, a digital representation of fibers as polylines, e.g. a variable sequence of points in 3D space. One of the most common purposes of tractography computation is the identification of the main white matter neuroanatomical pathways, namely the fiber bundles. The task of fiber bundle segmentation aims to select the portion of streamlines that belongs to a known white matter fascicle [2]. Several neuroimaging studies require comparing these brain white matter structures with respect to a template [3] or among a population [4], either healthy individuals or patients. In both cases, the mandatory premise is the registration of these data to a common space. While the registration of volumetric data can rely on a shared grid of voxels, the vector representation of streamlines poses a completely different problem, usually known as tractogram alignment. Differently from voxel representation, where we have a pairwise correspondence of voxels between the images of two individuals, we can’t have the similar relations among the 3D points that define the pathway of a streamline.

So far, the best practice of tractogram alignment is based on the application of the transformation computed from volumetric reference/related images [5]. The usual workflow is the following: (i) computation of the transformation from a *moving* to a *static* anatomical MRI T1-weighted (T1-w) image, (ii) application of the transformation to the tractogram of the moving individual. The tacit assumption is that tractography and the corresponding MRI images are already aligned with each other at the level of the same individual.

The general concern is that the transformation based on T1-w images is neglecting the more accurate information of white matter tissues encoded by DWI data. Several methods have been proposed to exploit a more accurate representation of the white matter by using DWI data derivatives. Considering the information at the voxel level, we have increasingly complex methods: fractional anisotropy (FA), diffusion tensor (DTI), and fiber orientation distribution (FOD) [1].

Despite the effort to improve the accuracy of registration using DWI data derivatives, the scientific literature is missing a comprehensive quantitative analysis of the impact and the benefits of these methods on tractogram alignment [5]. The major issue is represented by the lack of ground truth for what should be considered the correct alignment of two tractograms. To overcome this limitation, we designed an evaluation framework where we used manually segmented bundles as fiducial points to assess the correct registration between two individuals. The novel contribution of this work is an extensive empirical assessment of how the most prominent methods for white matter registration behave on a publicly available dataset.

## 2. RELATED WORKS

The most common approach to tractogram alignment is based on transformations computed on volumetric images. The transformation is then applied to the single 3D points of the streamlines [5]. The simplest option is to compute a linear or non-linear transformation taking in input the T1-w images, i.e. a gray intensity scalar data. In this case, the registration is mainly driven by the folding of gray matter because the information on white matter tissues is poor.

DWI acquisition enables the computation of data derivatives, such as diffusivity metrics. The most familiar is fractional anisotropy (FA), a scalar measure that indicates the mean voxel level of diffusion anisotropy of water in tissue. The FA emphasizes the contribution of high anisotropic tissue, i.e. the white matter fibers respect the low anisotropic brain regions, e.g. the brain ventricles.

The registration based on diffusion models, derived from DWI data processing such as DTI and FOD, presents a more challenging calculation of the transformation than using T1-w images. In addition, to estimate the spatial displacement for each voxel, we need to preserve information on the orientation of the diffusion, related to the direction of the fibers. The registration becomes an optimization process that combines voxel displacement and reorientation.

For DTI registration, there are two main approaches for the reorientation of the diffusion tensor: Finite Strain (FS) and Preservation of Principal Directions (PPD) [6]. In the former, the transformation matrix is decomposed in its rotation part and deformable part, and the rotation is applied to reorient the tensor at voxel level while in the latter the transformation is directly applied to the eigenvectors of the tensor. In PPD, there is the assumption that the main direction is encoded by the major eigenvector, therefore the reorientation of the other two eigenvectors is obtained preserving the orthogonality.

In the case of higher order tensor, the single fiber orientation assumption doesn’t hold anymore. The single rotation is not enough to reorient the FOD at the voxel level. In Preservation of Principal Branches (PPB) each main fiber orientation is rotated separately [7]. A more recent method is proposing to first approximate the FOD as a sum of weighted Point Spread Function (PSF), to estimate the registration at the PSF level and then obtain the FOD reorientation as a weighted sum of PSF reorientation [8].

An entirely different approach to tractogram alignment is the computation of the transformation directly from streamlines rather than volumetric DWI derivatives [9][10][11]. In Streamline-based Linear Registration (SLR) [11], after a sub-sampling of the tractogram to a small subset of streamlines using clustering, an affine transformation is computed that minimizes a cost function based on Bundle Minimum Distance (BMD) [11].

Although SLR method is driven by tractography, the approach is still aimed to estimate a transformation in 3D space. The output of this method is an affine matrix defined over the voxel grid of a reference MR image. The problem of tractogram alignment has been recently reformulated as finding the correspondence among streamlines. According to this perspective, different solutions have been proposed based on graph matching [12], linear assignment problem (LAP) [13], and optimal transport [14]. The output of these methods is no more a transformation, either linear or non-linear, but a pairwise relation that defines the correspondence between the streamlines of two tractograms of two distinct individuals.

Typically, linear registrations are adopted in studies aimed to preserve inter-individual differences, while non-linear transformations are pursued for example-based bundle segmentation. Inter-individual difference might be investigated using the morphometry analysis with non-lineare (e.g. diffeomorphic) transformations.

## 3. MATERIALS AND METHODS

While the scientific literature on white matter registration provides a wide range of solutions, little is known about how different methods may affect tractogram alignment. The lack of comparative studies is not surprising. A quantitative analysis would require a sample of ground truth, where the correct alignment between two tractograms is defined. For this reason, the previous works are limited to bundle alignment [11, 15] rather than whole brain tractogram, or an indirect evaluation of the tractography alignment [5], or just a qualitative evaluation of the tractogram after registration [10, 14].

Our proposal is to exploit the knowledge of neuroanatomical white matter pathways in the brain. After the segmentation of known fiber bundles, we can establish the correspondence between homologous structures as fiducial points for the assessment of tractogram alignment. According to this view, the requirement is the expert annotation of tractograms for a sample of individuals.

For this purpose, we consider TractoInferno^1^[16], an open publicly available dataset of diffusion MRI data. The collection includes DWI data and its derivatives (FA, DTI, FOD, tractography) and anatomical MRI data (T1-w) of 243 individuals acquired from 6 different sites with a 3T MR scanner. The added value of this dataset are the segmented bundles, 30 different ones, manually revised by a team of experts from an ensemble tractography of 13 millions of streamlines.

Unfortunately, the 30 bundles have not been segmented for each individual. The bundles identified with inaccurate anatomy were not included in the data publication. For this reason, we selected all the individuals where all the 30 bundles were available. Furthermore, we applied the additional constraint to include only individuals where the cerebellum was acquired entirely in the MRI images. Since only 4 individuals met these requirements, we relaxed to 29 bundles obtaining a sample of 9 individuals. With these premises we were enabled to carry out 72 pairwise registrations from the group of 9 subjects.

We aggregated the bundles in 4 different groups: 4 commissural (corpus callosum: frontal lobe anterior/posterior, occipital and parietal lobe), 8 projection (pyramidal tract L/R, parieto-occipito pontine tract, fronto-pontine tract L/R, optic radiation L/R), 16 association (arcuate fasciculus L/R, inferior fronto-occipital fasciculus L/R, cingulum L/R, uncinate fasciculus L/R, superior longitudinal fasciculus L/R, inferior longitudinal fasciculus L/R, middle longitudinal fascicle L/R, frontal aslant tract L/R), 1 cerebellar (middle cerebellar peduncle). The average number of streamlines for each bundle is in the range 20*K* − 60*K*, and for the whole brain of each individual around 1 million. In Figure 1 we report the coverage of the whole brain anatomy given by the selected bundles.

**Fig. 1.**
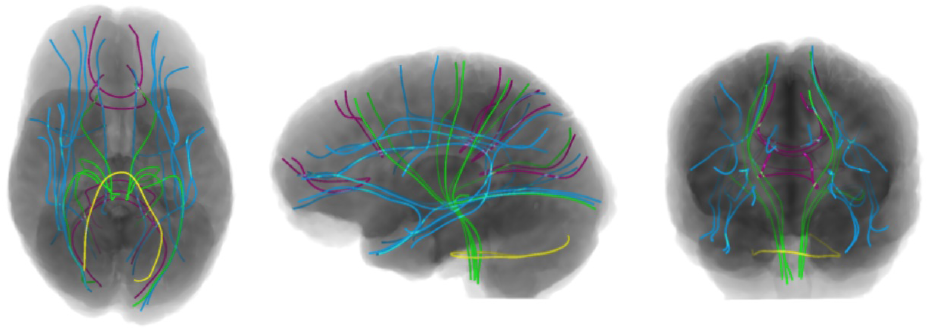
The visualization of one example of the main pathways for the 29 selected bundles for sub-1178 to support the pairwise evaluation of tractogram alignment: commissural (burgundy), projection (green), association (blue), cerebellar (yellow).

## 4. EMPIRICAL ANALYSIS

The aim of this study is an empirical assessment of a representative selection of methods for white matter registration to understand their performance on tractogram alignment. The ultimate goal is to provide some useful suggestions to orient the practitioners.

Given the huge number of methods proposed in the literature, we operated a selection following two main criteria: (i) the coverage of the main different approaches, and (ii) the availability of working implementation. As baseline registration, we computed a rigid transformation using ANTs [17] giving in input respectively T1-w and FA. For rigid registration based on tensor data, we referred to `DTIREG` of *DRTAMAS* software [18], part of the *TORTOISE* package [19], while for FOD registration [8] we referred to `mrregister` implemented on *MRtrix3* toolkit [20]. We computed the rigid transformation for streamlines using SLR [11]. In the next step, we considered methods for affine registration. We computed an affine transformation using ANTs taking in input T1-w and FA, respectively. For the same inputs, we computed the non-lineare registration as diffeomorphic transformations with ANTs. Affine and diffeomorphic transformation for DTI and FOD data were performed using `DTIREG` and `mrregister` respectively. For tractography based method, we computed the affine transformation with SLR while for correspondence methods we referred to LAP [13] because the implementation based on optimal transport does not scale up to 10^7^ streamlines.

The design of the empirical analysis follows a pairwise strategy: each method was applied to a pair of individuals, moving and static, respectively. As first step we computed the transformation from moving to static, the second step is the application of transformation to the moving tractogram, and the third step is the assessment of the mismatch between homologous bundles of moved and static tractograms. We repeated this procedure 36 times for all pairs drawn from the sample of 9 individuals, assuming symmetry of the registration.

We measured the evaluation of tractogram alignment at the level of bundle matching both for voxel representation and streamlines representation. In the first case, the bundle alignment is measured as the percentage of volume overlap by the Dice Similarity Coefficient (DSC). The volume of a bundle is obtained by computing the binary mask of voxels crossed by the streamlines. As a streamline-based measure, we refer to Bundle Minimum Distance (BMD) [11], the averaging of Minimum Direct Flip (MDF) distance among the streamlines of the two bundles.

We considered 4 image registration methods based on volumetric representations and 2 based on streamlines, 1 oriented to transformation and 1 to the criteria of correspondence. For the different kinds of input, we investigate separately the rigid, affine, and diffeomorphic transformations.

## 5. RESULTS AND DISCUSSION

In Figure 2 are reported the results of the empirical analyses to compare the different methods. The DSC scores of volumetric alignment of bundles are aggregated by the type of transformation (rigid, affine, and diffeomorphic) and by the category of brain pathways (commissural, projection, and association). Note that, the SLR method doesn’t support non-linear (e.g. diffeomorphic) transformation, while LAP doesn’t compute a transformation but the pairwise correspondence among streamlines. For the limit of space, we don’t report the scores of BMD, but the results are compliant with DSC.

**Fig. 2.**
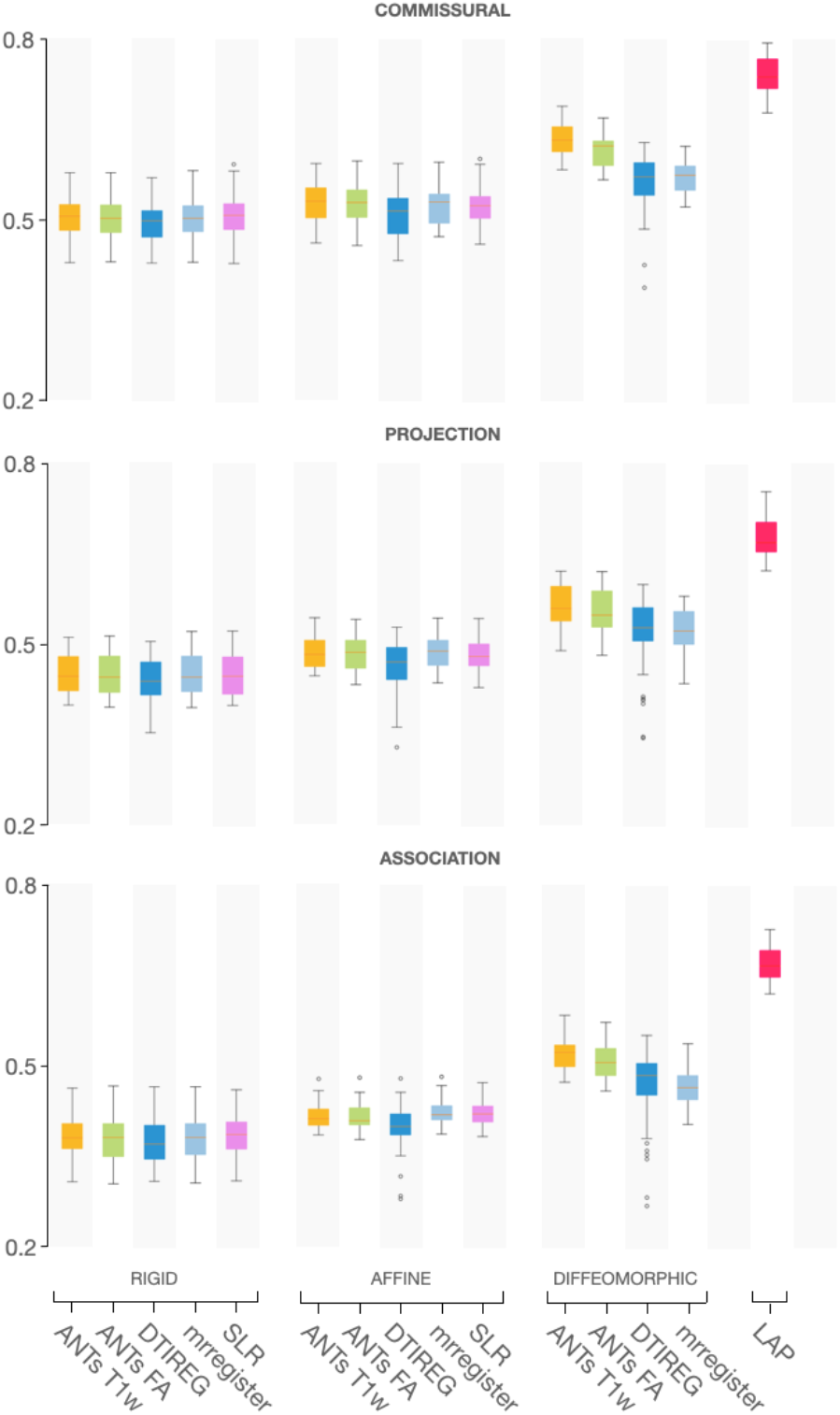
Volumetric Dice Similarity Coefficient for all registration methods aggregated as rigid, affine, and diffeomorphic transformations for commissural, projection, and association bundles respectively.

According to the expectation, a diffeomorphic transformation provides a boost in the alignment of tractograms, irrespective of the type of DWI data derivatives used for the registration process. The results for affine transformations are more consistent within the group than diffeomorphic ones, while the increase with respect to rigid transformations is marginal.

We may notice that the DSC scores tend to decrease across commissural, projection, and associative bundles, suggesting that the complexity of tractogram alignment is not uniformly distributed over the brain. Bundles where the length is longer and the shape irregularity is greater are more difficult to align.

Unexpectedly, the use of richer information on fiber orientation derived from diffusion models doesn’t seem to improve the alignment of tractograms. Single (DTI) and manifold (FOD) voxel-based fiber orientations don’t contribute to achieve a better registration of the streamlines. One explanation might be related to the spatial distribution of white matter, mainly in the inner part of the brain where the interindividual variance is lower. The registration of the outermost portion of the brain is dominated by the folding of the gray matter tissue, and the contribution of DWI data derivatives is negligible. Another option is the combination of manifold sources using a multichannel approach [5, 18]. Our trial using both T1-w and FA didn’t provide a meaningful difference.

The most newsworthy result is the gap of DSC score achieved by LAP. This outcome suggests that the criteria of streamlines correspondence might be more informative for tractogram alignment than voxel-based transformation. Further investigation is required to understand whether this difference might be explained by the underlining strategy. The streamline capture more comprehensive information about the fiber pathway rather than the voxel that encodes local orientation without the context of the whole pathway.

A deeper interpretation of results allowed us an additional finding that might be helpful in improving the tractogram alignment. In Figure 3 we report the usecase of alignment of the pyramidal tract between two individuals (sub-1160 and sub-1178). The pictures show the voxel mask of the bundle, the related density map of the streamlines, and the representation of the main pathway, e.g. the skeleton of the bundle [21, 22]. The skeleton of a bundle represents the mean pathway of the maximum density area of the streamlines, which does not necessarily coincide with the medoid of the bundle, e.g. the most representative streamline. We can visually have evidence that the voxel-based transformations are neglecting the information on fiber density.

**Fig. 3.**
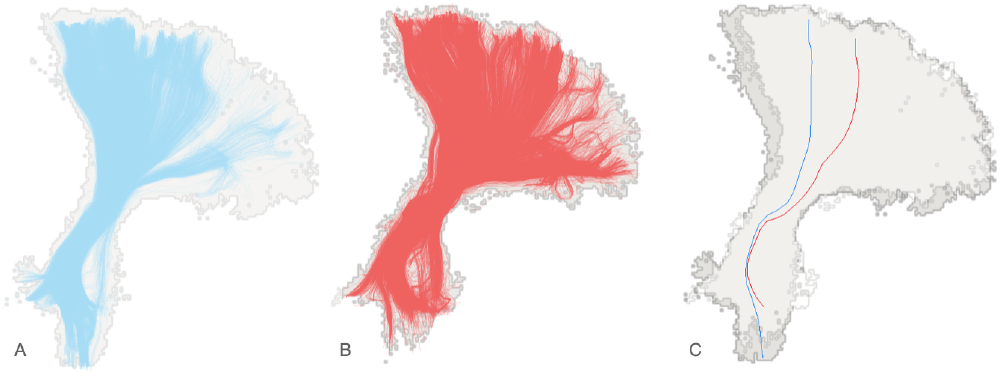
Example of bundle alignment: left pyramidal tract (PYT-L) between two individuals. (A) The voxel mask (gray) of the PYTE-L bundle of sub-1160 and the fiber density distribution (blue). (B) The voxel mask of PYT-L of sub-1178 warped to sub-1160 and the related fiber density distribution (red). (C) The overlap of the two voxel masks of PYT-L of both individuals and the skeleton of two bundles in the sub-1160 space.

Our finding is in accordance with previous studies [15, 23] on bundle alignment. A similar skeleton-like approach is based on Fiber-Flux Diffusion Density (FFDD) [15], where fiber density information is included in the registration process, which is limited to the alignment of the bundles separately. The open challenge is how to scale up from bundle to whole brain tractogram.

## 6. CONCLUSIONS

This empirical study shows how the state-of-the-art methods for white matter registration behave when the goal is the alignment of tractograms. We may conclude that even considering information on fiber orientation derived from diffusion models is not beneficial to the accuracy of fiber alignment between two individuals. What might be effective for voxel-based registration not necessarily has an impact on streamlines representation.

We found that registrations based on information of the whole streamline pathway are more effective than registrations based on local fiber orientation at the voxel level. The evaluation setup proposed in this work might be helpful to support further investigations on how to improve the tractography alignment.

## 7. COMPLIANCE WITH ETHICAL STANDARDS

This research study was conducted retrospectively using human subject data made available in open access at https://doi.org/10.18112/openneuro.ds003900.v1.1.1. Ethical approval was not required as confirmed by the license attached with the open access data.

## Footnotes

1 https://doi.org/10.18112/openneuro.ds003900.v1.1.1

## Notes

This work was partially supported by the grant PAT Reg. n. 764/2021 NeuSurPlan.

### Competing Interest Statement

This work was partially supported by the grant PAT Reg. n. 764/2021 NeuSurPlan.

